# Discovery of medicinal herbal compounds with potential anti-cancer activities against microtubule affinity-regulating kinase (MARK4) in cancer therapy

**DOI:** 10.1101/2023.05.30.542909

**Authors:** Nayana Narayanan, K.C Sivakumar

## Abstract

MARK4 belongs to the serine/threonine family and is found to be involved in apoptosis and many other regulatory pathways. Therefore, MARK4 is considered a potential target for cancer therapy. HTVS and XP of LOTUS and NPACT revealed that Ligand 11 and Ligand 7 respectively show good binding affinity along with ADME properties towards MARK 4. Further MD simulations for 50 ns suggested that the binding mechanism of Ligand 11 and 7 stabilizes the MARK4 by forming a stable complex. Both the ligands were bound to the active site of MARK4. This work provides a new insight into the use of Ligand 7 and Ligand 11, which were obtained from herbal extracts belonging to the class of Flavonoids and Megastigmanes, respectively, showing anticancer activities. The MD simulation studies suggest that Ligand 11 and Ligand 7 can be considered as potential inhibitors to MARK 4. Overall, this study provides an experimental evaluation of the herbal compounds identified during the study against MARK 4-associated cancers

## Introduction

MARK4 (microtubule affinity regulating kinase) is a potential therapeutic target for a number of cancers because it controls the initial stages of cell division (1). Every organ in the human body expresses the MARK4 gene, which is located on chromosome 19. The growth of gliomas, lung cancer, and leukemia are all influenced by its up-regulation (2). MARK4 was discovered as a result of its ability to phosphorylate tau proteins in microtubules of neurons and other proteins that are associated with microtubules (3). The development of prostate cancer is aided by MARK4, which is connected to -catenin/TCF-dependent transcription activity and is also a component of the Wnt signaling pathway (4). Recent reports that describe MARK4’s role in a number of diseases (5) have highlighted the significance of this protein. MARK4 overexpression has been associated with diabetes and obesity, AD, and metastatic breast cancer (4). Recent research (3) has demonstrated its role in the development of breast cancer metastatic spread. The highest levels of MARK4 expression can be found in the testes, kidney, and brain. In addition to NF-B, mTOR, Wnt, and Akt, MARK4 expression is linked to a number of signaling pathways and inhibits the Hippo signaling pathway, which encourages pro-liferation (3, 6, 7). As a result, MARK4 kinase is being developed as a primary target for therapeutic treatment of a number of diseases (2–4, 8, 9). MARK4 has a catalytic domain at the N-terminus, a linker loop, a UBA domain, a common docking motif, a C-terminal tail, and a spacer (10). MARK4S and MARK4L are splice variants of MARK4 that differ in the C-terminal region (8). MARK4S is extensively expressed in the brain and has a role in neuronal development, whereas MARK4L is largely expressed in the testis, where it is also found expressed in both sertoli and germ cells. They are also up-regulated in glioma cells, hepatocellular carcinomas, and metastatic breast carcinomas. However, studies show that MARK4L is linked to a decrease in -catenin/TCF-dependent transcription activity and that it can also act as a negative regulator of mTORC1. All these characteristics make MARK4L a viable target for cancer treatment (4). With the discovery of MARK4’s comprehensive properties, there has been a surge in efforts to find small compounds that can block MARK4, hence lowering the impacts of MARK4 upregulation and overexpression (4, 8, 11–13). All of this suggests that inhibiting MARK4 with small molecular inhibitor drugs lowers the phosphorylation of tau proteins and other microtubule-associated proteins, delaying the advancement of various abnormalities connected to upregulation or overexpression of MARK4 will be an excellent therapeutic strategy against diverse types of cancers and associated diseases (14, 15).

The proposed theories of MARK4 as a potential target for anticancer drug development provide a way for researchers to discover new compounds. Many of the MARK4 inhibitors discovered so far are still in the preclinical stage (16). According to a study (17), PCC0208017, a synthetic drug that showed excellent antitumor action against glioma during molecular docking experiments, had a high binding affinity for MARK3 and MARK4, and was discovered to be a new inhibitor of both MARK4 and MARK3. AChE inhibitors, such as Donepezil and Rivastigmine Tartrate, have been found to inhibit MARK4, providing a treatment option for Alzheimer’s disease (18). Another study performed 59 pyrazolopyrimidine derivatives in-silico to determine their affinity and molecular docking with MARK4 and found five of them to be potential inhibitors (19).

A series of arylaldoxime/5-nitroimidazole hybrids (4a to 4h) were developed and synthesized in the quest for effective and specific MARK4 inhibitors, and 4h was discovered to be a robust inhibitor of MARK4 kinase (13). Researchers synthesized isatin-triazole hydrazones in the search for an effective inhibitor (9a to 9i). Following docking studies, it was discovered that 9g has a higher affinity for MARK4, which aids in cell growth control and the addition of an extra repository for cancer therapeutic medicines. (12). Another study involved synthesizing and designing small chemical inhibitors of MARK4 kinase using hydroxylamine derivatives of morpholine. According to molecular docking experiments, compound 32 (N- (morpholin-4-yl)[2-(trifluoromethyl)phenyl] methylidenehydroxylamine) has a higher affinity for MARK4 than other produced compounds. (20). Many investigations supported the idea of replacing synthetic drugs with natural compounds that demonstrated less toxicity in a cytotoxicity experiment as a MARK4 inhibitor. Rosmarinic acid is a naturally occurring phenolic chemical that has a high affinity for MARK4. (1)It is obtained from the Lamiaceae (mint family) family of plants. Another chemical, naringenin, which belongs to the flavanone glycoside group of polyphenols and is present in citrus fruits, has been identified as a possible inhibitor of MARK4 and could be used to treat cancer and other disorders caused by MARK4 expression. (21). The therapeutic properties of dietary flavonoids and polyphenolic substances suggest that MARK4 could be inhibited. As a result of the molecular docking experiments, two natural chemicals, rutin and vanillin, were discovered to be promising inhibitors of MARK4 (22). Citral is a pharmacologically active open-chain monoterpenoid isolated from lemongrass, and it has a significant affinity for MARK4. (23). -Mangostin is a dietary xanthone that has been reported to be a powerful MARK 4 inhibitor with a moderate capacity to slow cancer growth (24). Because bile acids are implicated in cancer-related processes such as the control of EFGR, cholic acid, an amphiphilic steroidal molecule, has been used to treat bile acid production disorders. According to comprehensive insilico research (25), cholic acid showed great binding affinity to MARK4 and These studies highlight the wide spectrum of biological chemicals found in nature that can be exploited as cancer and other illness therapeutics. According to the International Agency for Research on Cancer, there were 1.3 million new cancer diagnoses and 10.0 million cancer deaths globally in 2020 (26). Breast cancer, lung cancer, and prostate cancer are the most commonly diagnosed cancers worldwide. It is predicted that by 2040, there will be 28.4 million cases, up 47 percent from 2020 (27). On the other hand, 4 the growing rate of cancer occurrence and mortality, on the other hand, has become a global health issue. So many therapeutic approaches are available to reduce the risk of cancer malignancy, including chemotherapy, radiation, immunotherapy, surgery, bone marrow transplantation, chemically derived drugs, etc. (28). These therapies might induce stress in patients, which can lead to health problems as a result of the medicines’ negative effects (29, 30). As a result, alternative treatments and therapies are becoming more popular. The scientific community is looking for new revolutionary small molecule medications that have fewer side effects and operate only on cancer cells, which has led to substantial study into the identification of plant-derived drugs generated from chemicals taken from plants with anticancer potential. The quest for anticancer medicines derived from plants began in the early 1950s and continues to this day (31, 32). Many bioactive chemicals that are effective for treating various abnormalities have been identified from plants and are currently being tested in clinical or preclinical research. According to several studies, over 60medicinal plants (28, 32). Plant-derived drugs have been proven to be better targeted at specific organs or systems in the human body. In comparison to manufactured or chemical drugs, plant-derived drugs work slowly and have no side effects and features that contribute to the popularity of plant-based drugs (33). Researchers have set a new target of developing anticancer medicines from herbal components with great selectivity and potency. The screening of natural compounds based on their chemical and structural variety from existing databases, examining their anticancer capabilities, and then going through binding affinity testing, etc. are the first steps in drug discovery (1). The objective of this research is to find an anticancer herbal component that has a high affinity for MARK4. Molecular docking studies will be used to obtain structural insights into the interaction between plant-derived compounds and MARK4, and binding affinities will be analyzed using molecular dynamic simulations.

## METHODS AND METHODOLOGY

This research was conducted on an NVIDIA DGX station A100 using a server-grade CPU with a single AMD 7742, 128 cores, and an Ubuntu Linux operating system at speeds ranging from 2.25 GHz (base) to 3.4 GHz (max boost) (20.04.) The software packages used for this study are part of a computing environment called Maestro by Schrodinger (version: 2021-1). For simulations, DESMOND was used, GLIDE was used for docking, and Qikpro was used to examine the ligands’ ADME properties. There are several steps to be done before high throughput virtual screening and docking using GLIDE

1. Receptor Grid Generation
2. Ligand Preparation
3. Glide Docking(screening)
4. PROTEIN PREPARATION

### Preparation of the Ligand Library and Protein

The crystal structure of human MARK4 with a resolution of 2.8 was retrieved from the Protein Data Bank (PDB ID: 5ES1) and used as the receptor (15). A database containing 2,765,518 natural compounds for the ligands was retrieved in a single SDF file format from the LOTUS database, and 504 herbal compounds having anticancer properties from the NPACT database were used for structural-based virtual screening. The PDB structure we obtained from various sources may lack the necessary information needed for performing modeling, like missing hydrogens, partial charges, sidechains, or whole loop regions. The protein preparation wizard is used to correct these deformities. In the present study, the 5ES1.PDB structure is used for the preprocessing and optimization process. The initial process in the import and process tab is filled with missing side chains/loops, removing water beyond 5 (depending on the needs). The refine tab imparts more modifications to the PDB structure, like optimizing the hydrogen bonding network.

#### 1. RECEPTOR GRID GENERATION

For glide docking, the initial step is to create a protein grid that represents the precalculated interaction properties of the protein. This grid generation will speed up the ligand docking step. The grid is generated as an expected binding point (usually by extrapolating the ligand-bounded ligand bounded region in the crystallographic structure). There are many advanced options for adding more information to the receptor grid. The grid generation usually takes 2-3 minutes to create a zip file, which is the prerequisite for ligand docking. A receptor grid was generated as the center coordinates (X=- 44.17, Y=-23.14, and Z=-4.25) with a ligand midpoint diameter box of dimension 10 × 10 × 10. The grid box was positioned at the centroid of the workspace ligand. If the structure in the workspace is a combination of receptor and ligand, the ligand should be selected to be excluded from the receptor grid generation.

#### 2. LIGAND PREPARATION

This is done by ligprep, a collection of tools for converting molecules. Ligprep can convert ligands to 3D structures readily with standardized and extrapolated. These conversions are done by applying corrections to structures by generating variations, eliminating unwanted structures, and optimizing them by generating tautomer and ionization states, making them ready for virtual screening. In this study, 276,518 compounds were from the LOTUS database, and 504 compounds showing anticancer activities were taken from the NPACT database.

#### 3. GLIDE DOCKING

The common computational strategy of structural-based ligand docking involves screening the compounds. Glide docks the flexible ligands to the rigid protein using rapid sampling of orientational and positional degrees of freedom of the ligand. There are three modes of Glide screening and docking. These three modes of Glide docking produce a set of conformers for a ligand and employ a series of hierarchical filters to enable rapid evaluation of ligand degrees of freedom.

#### 4. HIGH THROUGHPUT VIRTUAL SCREENING

##### 4.1 STANDARD PRECISION SP

is also a method of screening compounds that is much slower than HTVS, so it applies to a smaller number of compounds. The SP glide will be applicable to the screened compounds after the HTVS glide (34).

##### 4.2 EXTRA PRECISION

Extra-precision docking and scoring employ a harder scoring function that was optimized to minimize the number of false positives in screening. XP is more precise and computationally intensive than standard precision. During the docking process, the Glide scoring function (G-score) was used to select the best conformation for each ligand. The 20 ligands with the best docking scores were chosen from both databases (34).

### ASSESSMENT OF DRUGABILITY

Assessment of druglike properties of selected ligands using the Qikpro suite for 20 compounds that are selected after screening. QikProp predicts significant descriptors and pharmaceutically applicable features of organic molecules, either individually or in batches. QikProp provides ranges for comparing a particular molecule’s properties with those of 95in this study, the compounds with zero-star value (#star) and good docking (Glide) scores were chosen for molecular dynamics simulation (35).

### MOLECULAR DYNAMICS SIMULATIONS

A MD simulation mimics the changes in the structures of biological molecules over a given period, giving us atomic insights into the change in structure and giving us detailed information about the fluctuations and conformational changes of the proteins and other small molecules under study (36). In this study, Desmond, an advanced MD simulation system having a graphical user interface integrated with Maestro, was used. In Desmond, we can perform simulations of small molecules, proteins, nucleic acids, and membrane systems to study the interactions of complex molecular assemblies. Desmond trajectories can be visualized and analyzed using numerous tools in Maestro. The system should be built using the system builder before moving to simulation. The System builder is the GUI that generates a solvated system for simulation. The solvated system-generated includes the solute (protein, protein complex, protein-ligand complex, or similar systems, etc.), solvent, and counter ions. All structural topological information and force field parameters for the solvated system are written to a special Maestro file that is subsequently used for Desmond simulation. The solvent model used for solvation is TIP4P. The solvated system was within a 10 box edge. The force field used is OPLS4. Cl counter ions were added for system neutralization, and MD simulation was run for 50 ns for selected candidates. The NPT ensemble class with a temperature of 300K and a pressure of 1 bar was applied for both MD runs.

## RESULTS AND DISCUSSION

### Screening of Lead Compounds

The HTVS method of Glide was used to screen the entire LOTUS and 504-specific compound databases from the NPACT database, and Glide extra precision was then used to dock the screened compounds (XP). Each low-energy conformer in the chosen binding site has its reasonable conformations determined by the glide XP mode (37). The first 20 compounds from HTVS screening were initially eliminated with a cutoff of -7.43Kcal/mol and subjected to Glide XP docking. Then, using Qikprop Schrödinger Release 2021-4, the 20 compounds with the highest binding affinities were permitted to be studied for their drug-like properties according to Lipinski’s rule of five. (35). Both the 20 compounds from the NPACT database with an estimated Glide XP docking score of-12.927 to-5.857 Kcal/mol and the 20 compounds from the LOTUS database with an estimated Glide XP docking score of-8.44 Kcal/mol to-14.185 Kcal/mol had a good binding affinity for MARK4 according to the Glide XP docking analysis table1 and table2. Additionally, by examining their phytochemical properties and identifying compounds with a star value of 0, the compounds’ drug-likeness was assessed using Qikprop in both databases table4 and table5. Based on the docking score and star value, Ligand-11 from the LOTUS database and Ligand 7 from the NPACT database were chosen for additional molecular dynamics simulation studies.

**Table 1.** Selected compounds from screening and their docking score after XP from LOTUS database.

| Compound No | Compound ID | Docking Score (Kcal/mol) |
| --- | --- | --- |
| Ligand 1 | LTS0005188 | -14.185 |
| Ligand 2 | LTS0003554 | -13.992 |
| Ligand 3 | LTS0000770 | -13.08 |
| Ligand 4 | LTS0003473 | -11.701 |
| Ligand 5 | LTS0004685 | -10.746 |
| Ligand 6 | LTS0004349 | -10.515 |
| Ligand 7 | LTS0000669 | -10.264 |
| Ligand 8 | LTS0001422 | -10.216 |
| Ligand 9 | LTS0004168 | -10.135 |
| Ligand 10 | LTS0003674 | -10.072 |
| Ligand 11 | LTS0005218 | -9.82 |
| Ligand 12 | LTS0003539 | -9.796 |
| Ligand 13 | LTS0003183 | -9.726 |
| Ligand 14 | LTS0000608 | -9.714 |
| Ligand 15 | LTS0000911 | -9.658 |
| Ligand 16 | LTS0001454 | -9.477 |
| Ligand 17 | LTS0005397 | -9.082 |
| Ligand 18 | LTS0005466 | -8.6 |
| Ligand 19 | LTS0002900 | -8.474 |
| Ligand 20 | LTS0005526 | -8.44 |

**Table 2.** Selected compounds from screening and their docking score after XP from NPACT database

| Compound No | Compound ID | Docking Score (Kcal/mol) |
| --- | --- | --- |
| Ligand 1 | 250395 | -14.185 |
| Ligand 2 | 6474309 | -13.992 |
| Ligand 3 | 10455035 | -13.08 |
| Ligand 4 | 6440783 | -11.701 |
| Ligand 5 | 475263 | -10.746 |
| Ligand 6 | 10895435 | -10.515 |
| Ligand 7 | 10498462 | -10.264 |
| Ligand 8 | 1638 | -10.216 |
| Ligand 9 | 10228895 | -10.135 |
| Ligand 10 | 14218027 | -10.072 |
| Ligand 11 | 65084 | -9.82 |
| Ligand 12 | 10722779 | -9.796 |
| Ligand 13 | 3071 | -9.726 |
| Ligand 14 | 6773 | -9.714 |
| Ligand 15 | 6444206 | -9.658 |
| Ligand 16 | 10585660 | -9.477 |
| Ligand 17 | 44575506 | -9.082 |
| Ligand 18 | 19889 | -8.6 |
| Ligand 19 | 134813472 | -8.474 |
| Ligand 20 | 9986592 | -8.44 |

### Drug likeness of lead compounds

The drug-like properties in accordance with Lipinski’s rule of five of the selected ligands exhibited the five properties are within the acceptable range; The drug-like behavior analysis of these two compounds was studied by analyzing the pharmacokinetic parameters required for absorption, distribution, metabolism, excretion, and toxicity (ADMET) by the use of QikProp table3. The partition coefficient (QPlogPo/w) and water solubility (QPlogS) are critical for the estimation of absorption and distribution of drugs within the body (37). Ranging between the recommended values, cell permeability (QPP Caco), a key factor governing drug metabolism and its access to biological membranes, is in an acceptable range. Overall, all the descriptors fulfill the drug-likeness properties with a Qikprop star value of zero for both, thereby indicating that they can be considered as potential drug candidates.

**Table 3.** Showing the Qikpro properties of identified ligand

| Compound<sup>a</sup> | QLogPo/wd<sup>b</sup> | QLogS<sup>c</sup> | QPP Caco<sup>d</sup> | QLogHERG<sup>e</sup> | % human oral absorption<sup>f</sup> |
| --- | --- | --- | --- | --- | --- |
| Ligand 11 | -0.577 | -2.033 | 55.244 | -4.045 | 54.748 |
| Ligand 7 | 2.706 | -4.516 | 136.531 | -5.059 | 81.009 |

<sup>a</sup>Identified compounds
<sup>b</sup>Predicted octanol/water partition coefficient log p (acceptable range: -2.0 to 6.5).
<sup>c</sup>Predicted aqueous solubility; S in mol/L (acceptable range: -6.5 to 0.5)
<sup>d</sup>Predicted Caco-2 cell permeability in nm/s (acceptable range: <25 is poor and >500 is great)
<sup>e</sup>Predicted IC 50 value for blockage of HERG K<sup>+</sup> channels (acceptable range: below -5.0)
<sup>f</sup>Percentage of human oral absorption (<25% is poor and >80% is high)

**Table 4.** #Star value after Qikprop analysis of the docked compounds from LOTUS database

| Ligands | Ligand ID | Docking Score (Kcal/mol) | #Starvalue |
| --- | --- | --- | --- |
| ligand 1 | LTS0005188 | -14.185 | 16 |
| ligand 2 | LTS0003554 | -13.992 | 14 |
| ligand 3 | LTS0000770 | -13.08 | 12 |
| ligand 4 | LTS0003473 | -11.701 | 4 |
| ligand 5 | LTS0004685 | -10.746 | 5 |
| ligand 6 | LTS0004349 | -10.515 | 3 |
| ligand 7 | LTS0000669 | -10.264 | 11 |
| ligand 8 | LTS0001422 | -10.216 | 9 |
| ligand 9 | LTS0004168 | -10.135 | 15 |
| ligand 10 | LTS0003674 | -10.072 | 3 |
| ligand 11 | LTS0005218 | -9.82 | 0 |
| ligand 12 | LTS0003539 | -9.796 | 4 |
| ligand 13 | LTS0003183 | -9.726 | 0 |
| ligand 14 | LTS0000608 | -9.714 | 0 |
| ligand 15 | LTS0000911 | -9.658 | 2 |
| ligand 16 | LTS0001454 | -9.477 | 2 |
| ligand 17 | LTS0005397 | -9.082 | 0 |
| ligand 18 | LTS0005466 | -8.6 | 0 |
| ligand 19 | LTS0002900 | -8.474 | 1 |
| ligand 20 | LTS0005526 | -8.44 | 0 |

**Table 5.**
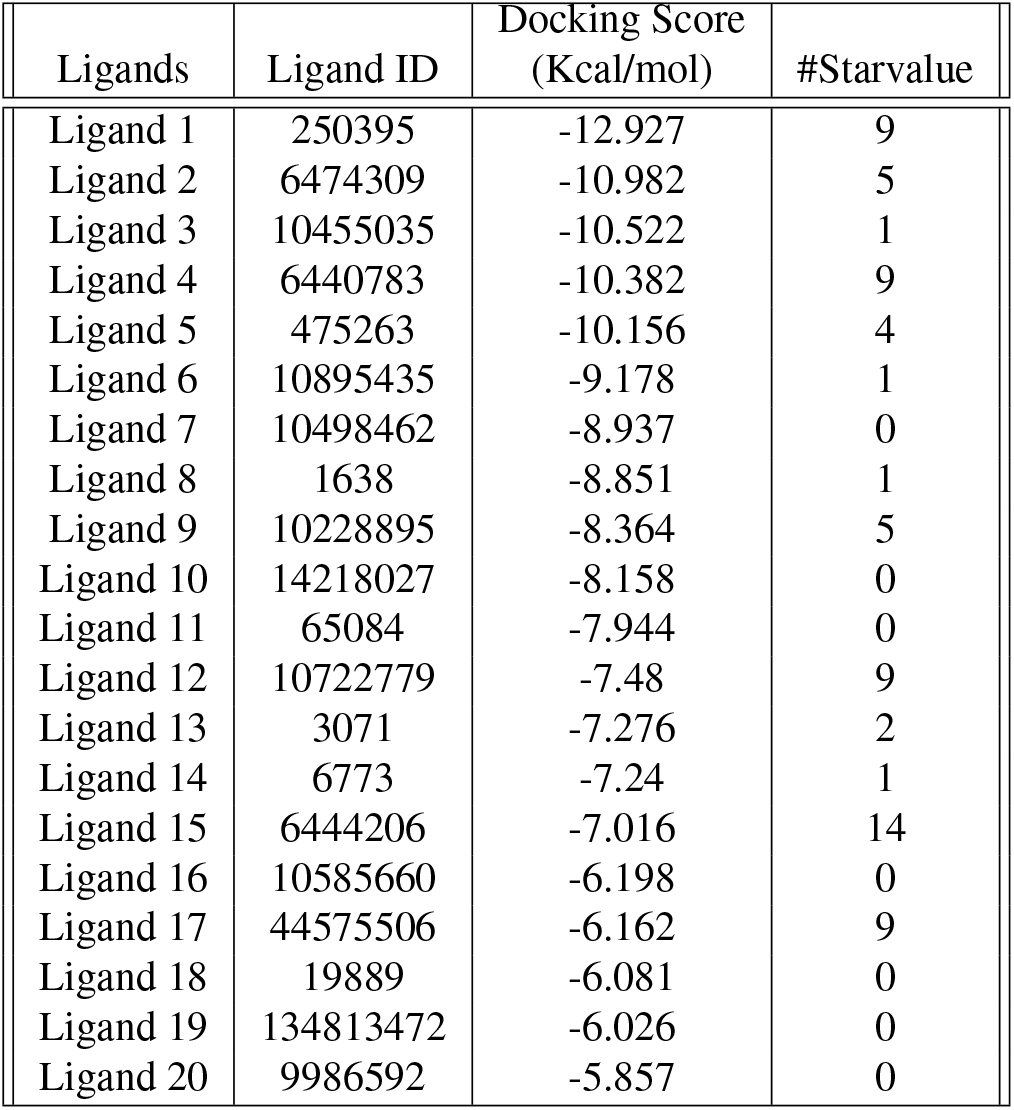
#Star value after Qikprop analysis of the docked compounds from NPACT database

### Molecular docking Analysis

The ligand-receptor complex is predicted using molecular docking. Ligands 11 and 7’s docked conformation with MARK4 sheds light on the binding interactions. figure1 and figure2 show that the orientation of the ligands in the binding pocket of MARK4 has been optimised by reducing the total energy. The significant binding affinity scores for the Ligand 11-MARK4 complex and 10 Ligand 7-MARK4 complex were -9.82 Kcal/mol and - 8.937 Kcal/mol, respectively. The ligand interaction diagram reveals that they interact hydrophobically and exhibit stable confirmation with the amino acids in the active site of MARK 4’s negatively charged amino acid residues. As shown in figure2, Ligands 11 and 7 formed hydrogen bonds with the amino acids GLU 139, GLU 133, and ASP 142 and ALA 135 and ASP 142, respectively, to bind to the MARK4 protein figure4. This docking study demonstrated the attachment of two chosen ligands to the previously identified binding cavity of MARK4, which is where its substrate binds. The investigated substances can act as inhibitors by lowering MARK4’s affinity for its substrate

**Fig. 1.**
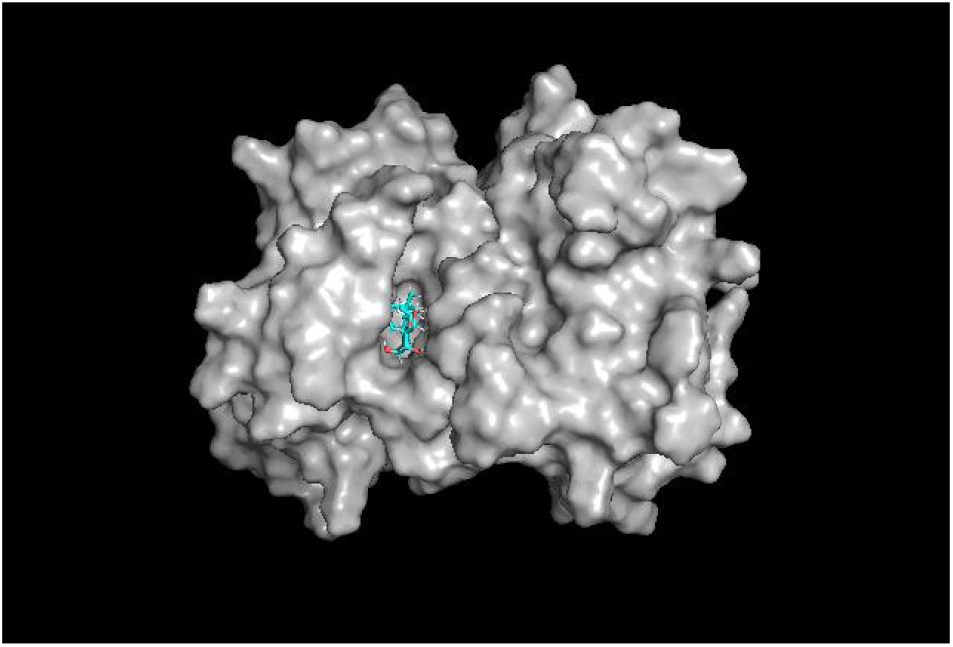
Image of Binding pocket of MARK4 along with Ligand 11

**Fig. 2.**
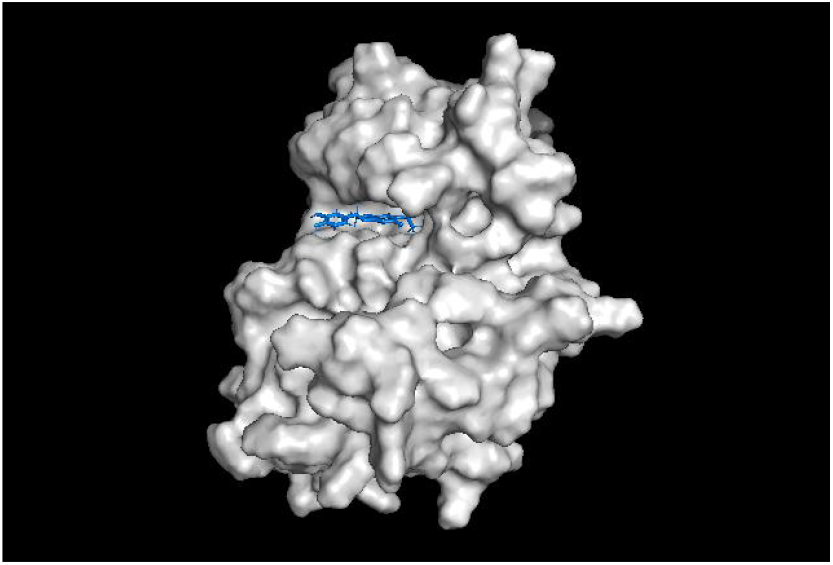
Image of Binding pocket of MARK4 with ligand 7

**Fig. 3.**
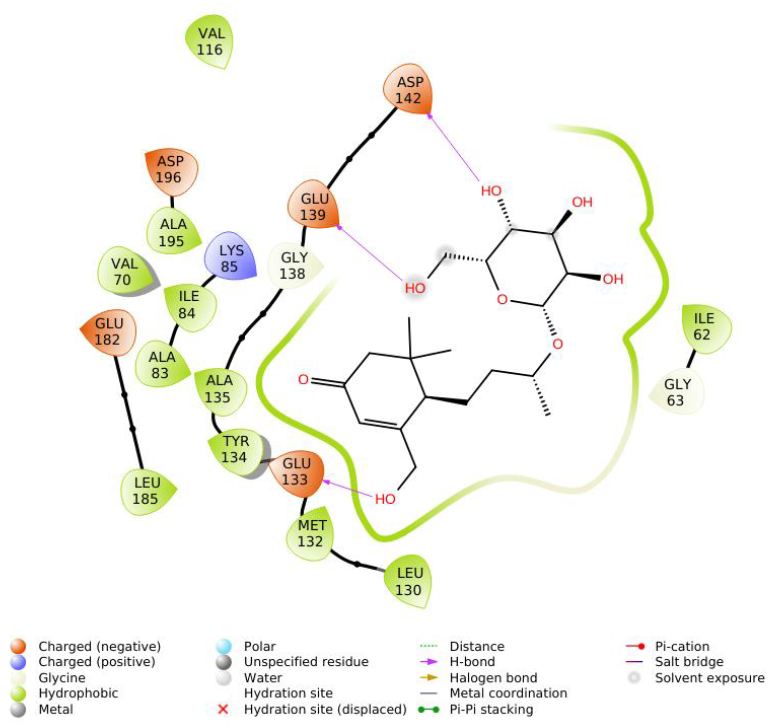
Interaction diagram showing the residues of MARK4 - Ligand 11 showing the residues of active site

**Fig. 4.**
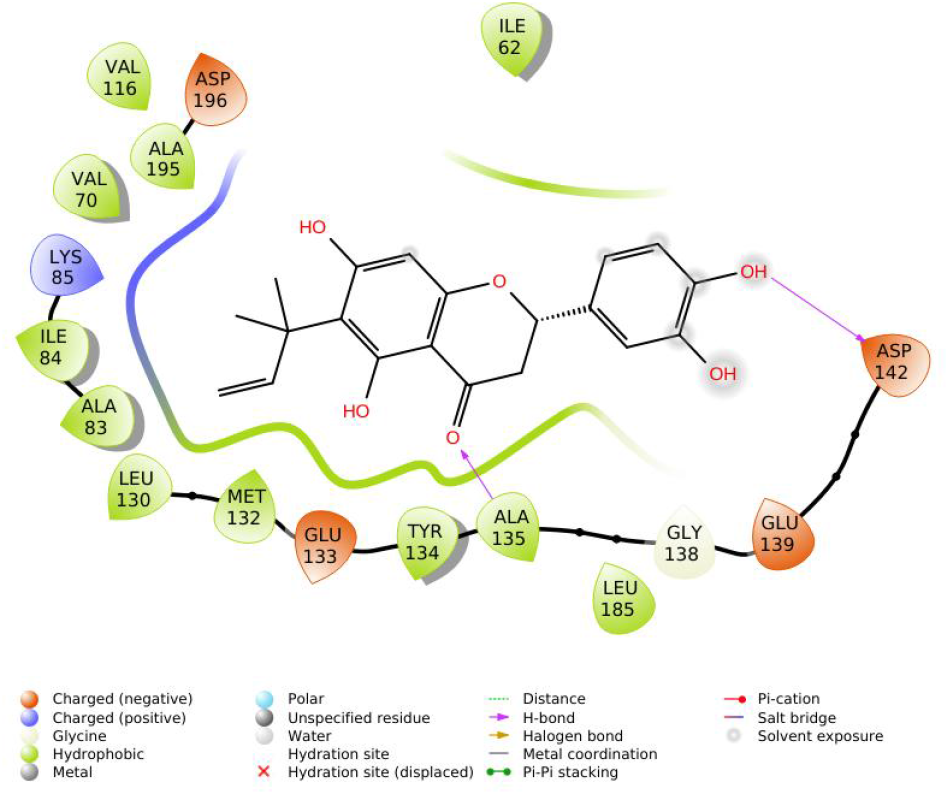
Interaction diagram of Ligand 7 with the residues of MARK4

### Molecular Dynamics Simulation

According to the molecular docking studies used in this study, Ligand-11 and Ligand-7 are assumed to have a close binding conformation. Schrodinger ran a molecular dynamics simulation for both ligands in the Desmond package for 50 ns in order to learn more about the interaction mechanism (36). The interaction between protein and ligand was investigated using this trajectory file, which was created for 4.8 ps intervals. By examining the RMSD, it was possible to keep track of the molecular dynamics of the protein-ligand complex. One of the fundamental characteristics used to assess the stability of protein structures is the root mean square deviation (RMSD)(38). The RMSD of unbound MARK 4 run for 50 ns is shown in figure5a, demonstrating how the protein’s structural integrity was maintained throughout the simulation. The plots in figure5c and figure5. display the RMSD of the MARK4-Ligand 11 and MARK4-Ligand 7 complexes for the 50 ns MD run, respectively. Comparing the graphs reveals that the MARK4 has been stabilised by the binding compounds, resulting in fewer structural variations. To find the average fluctuations of all residues during the simulation process, the root mean square fluctuations (RMSF) of MARK4 were additionally plotted figure6a. Additionally, the RMSF of bound MARK4 was plotted, showing that fluctuations were reduced during compound binding figure6b and figure6c. It was discovered that Ligand 11 binding exhibits greater fluctuations than Ligand 7.

**Fig. 5.**
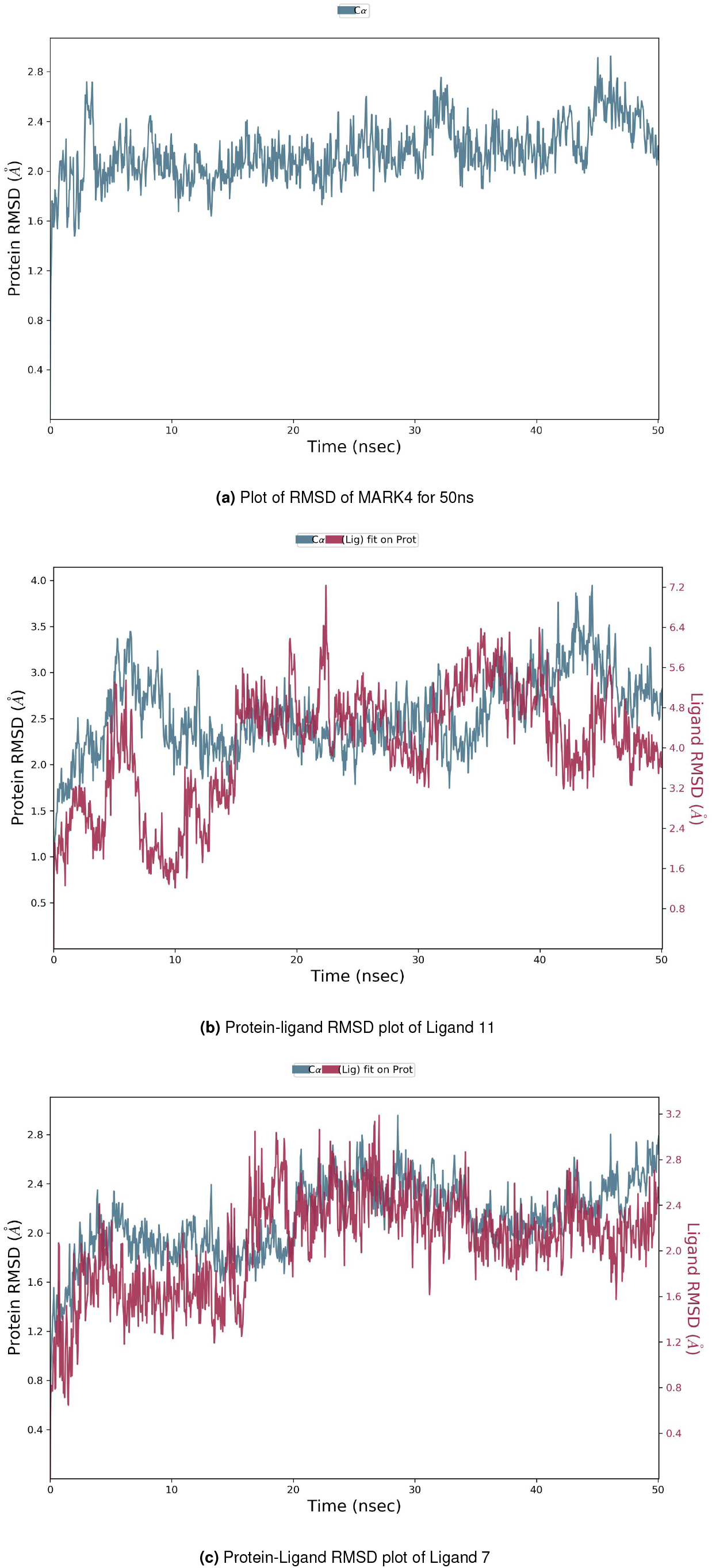
RMSD

**Fig. 6.**
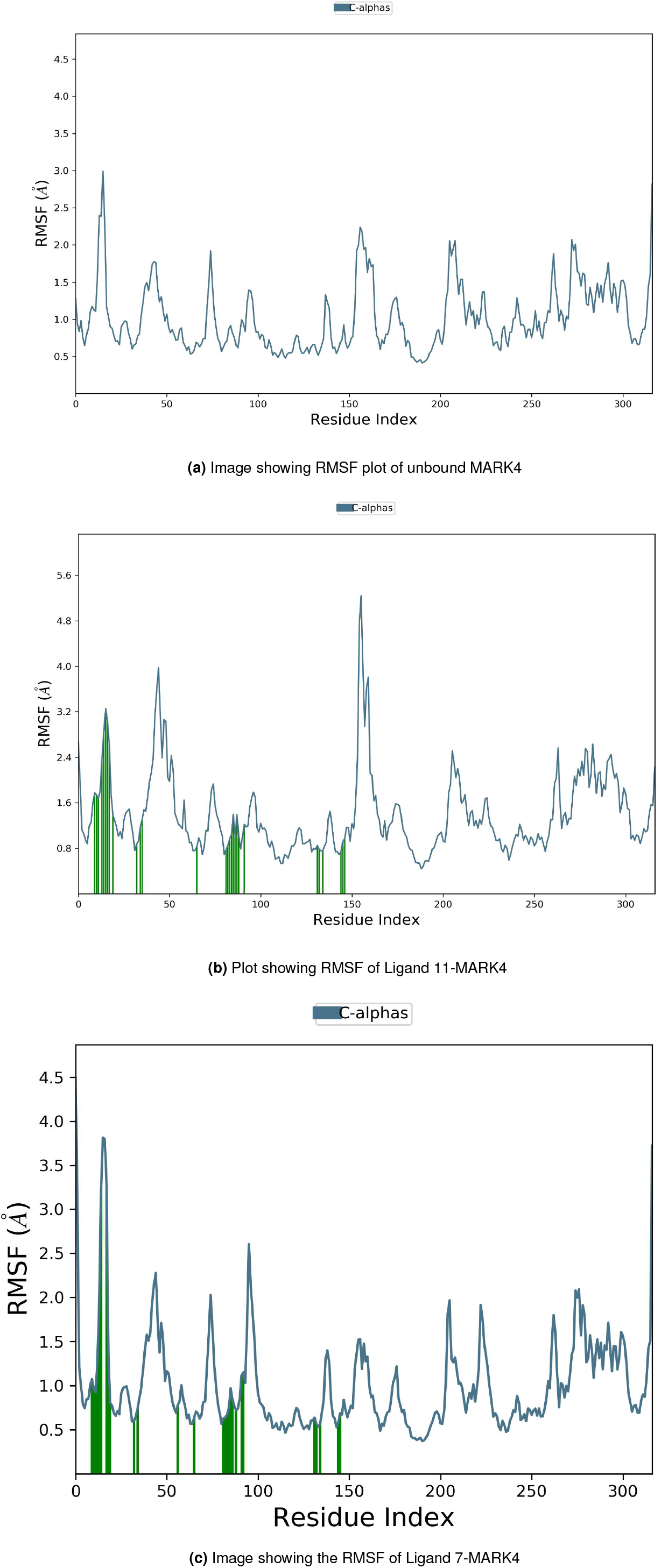
RMSF

### A. Protein-Ligand Contacts

Throughout the simulations, protein interactions with ligands were observed. Hydrogen bonds, hydrophobic, ionic, and water bridge interactions make up these interactions. The analysis revealed that Ligands 11 and 7 had three to four hydrogen bonds holding them to the pocket of MARK4 (Figure7). While ALA 135 provided a hydrogen bond interaction with the ligand in ligand 7, GLU 133 demonstrated one with the ligand in ligand 11. Both ligands were interacting hydrophobically as well as through hydrogen bonds with the residues. The ligand binding modes and docking calculations were very similar. Overall MD simulation studies supported the theory that both ligands could serve as MARK 4 targets.

**Fig. 7.**
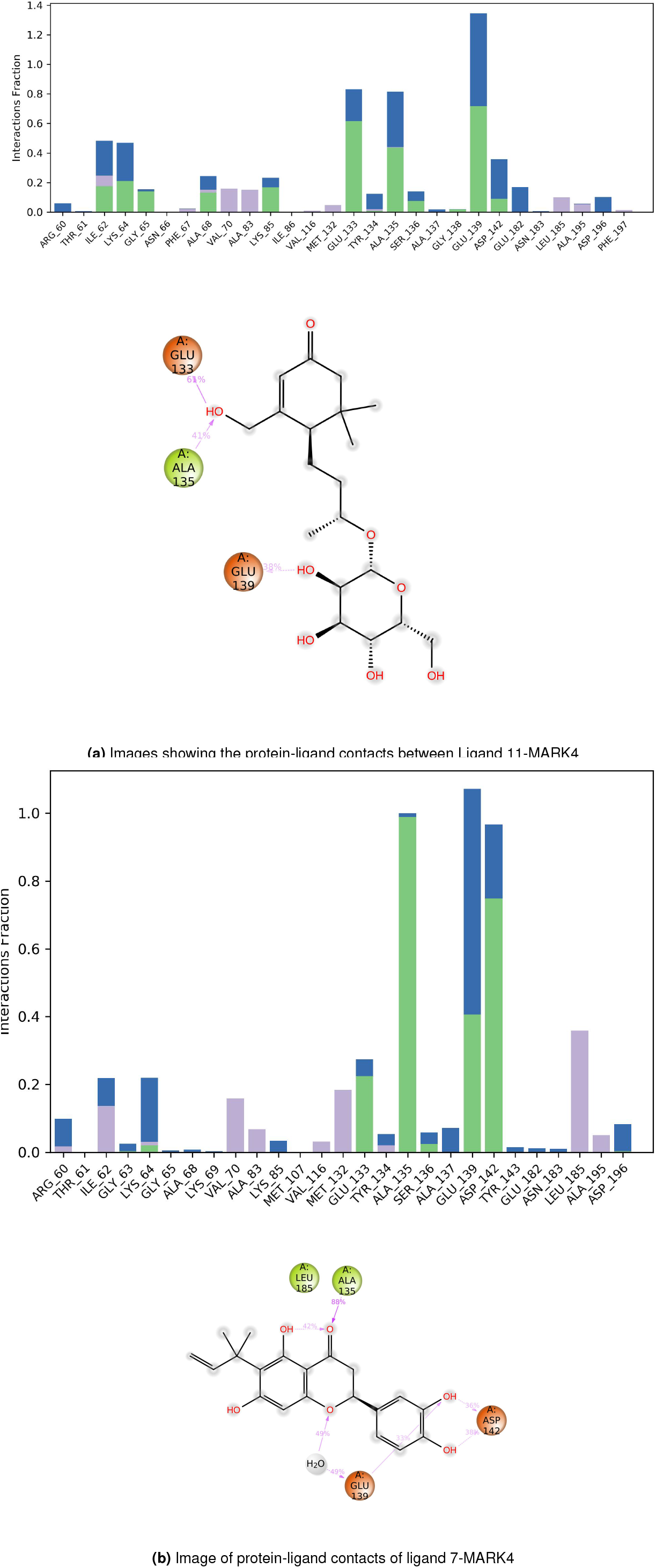
Protein-Ligand Contacts

### B. Lead compounds Information

The compounds identified belong to the class of Megastigmanes and flavonoids. Ligand 11 belongs to the class Megastigmanes and Ligand 7 belongs to the class flavonoids. The IUPAC name of Ligand 11 is 3-(hydroxymethyl)-5,5-dimethyl-4-(3-[3,4,5-trihydroxy-6-(hydroxy methyl)oxan-2-yl]oxybutyl)cyclohex-2-en-1-one and Ligand 7 is 6-(1,1-Dimethylallyl) eriodictyol. Both the compounds are from plant extracts, where ligand 11 can be obtained from four different plant species. The extracts from these plants are already in use for anticancer activities: Ornithogalum umbellatum (39), Allium obliquum, Salacia chinensis (40), Macaranga tanarius. The Megastigmanes were already identified for their anti-proliferative, anti-cancer, and cytotoxic effects. Most of the plants belong to the Indian habitat, except Allium obliquum. Ligand 7 belongs to the class of flavanones. It comes under the superclass of flavonoids, which possess several medicinal benefits, like antioxidative, anti-inflammatory, anti-mutagenic, and anti-carcinogenic properties, and the capacity to modulate key cellular enzyme functions. Ligand7 is extracted from the plant Monotes engleri and has medicinal value, being used to treat many cancer cell lines.

## CONCLUSION

The development of inhibitor compounds for kinases has become a promising approach to fighting against various diseases. This study concentrates on the identification of an inhibitor compound that is available in nature against MARK4 kinases. Screening of two different natural databases revealed that there are various herbal compounds that show good binding affinity towards MARK4. Since natural compounds show medicinal values, replacing synthetic inhibitors with natural ones will reduce the side effects of therapies. Studies have shown that flavonoids can bind with MARK4 better and Ligand 7, being a flavonoid compound, can act as a better inhibitor of MARK4. Although Ligand11 also shows anticancer activities, this study adds new compounds for therapeutic purposes, which further needs to be validated.

## REFERENCES

1. Saleha Anwar, Anas Shamsi, Mohd Shahbaaz, Aarfa Queen, Parvez Khan, Gulam Mustafa Hasan, Asimul Islam, Mohamed F Alajmi, Afzal Hussain, Faizan Ahmad, et al. Rosmarinic acid exhibits anticancer effects via mark4 inhibition. Scientific Reports, 10(1):1–13, 2020.

2. Farha Naz, Farah Anjum, Asimul Islam, Faizan Ahmad, and Md Imtaiyaz Hassan. Microtubule affinity-regulating kinase 4: structure, function, and regulation. Cell biochemistry and biophysics, 67:485–499, 2013.

3. Emad Heidary Arash, Ahmed Shiban, Siyuan Song, and Liliana Attisano. Mark4 inhibits hippo signaling to promote proliferation and migration of breast cancer cells. EMBO reports, 18(3):420–436, 2017.

4. Pranitha Jenardhanan, Jayakanthan Mannu, and Premendu P Mathur. The structural analysis of mark4 and the exploration of specific inhibitors for the mark family: a computational approach to obstruct the role of mark4 in prostate cancer progression. Molecular bioSystems, 10(7):1845–1868, 2014.

5. Alessandro Beghini, Ivana Magnani, Gaia Roversi, Tiziana Piepoli, Simona Di Terlizzi, Ramona F Moroni, Bianca Pollo, Anna M Fuhrman Conti, John K Cowell, Gaetano Finocchiaro, et al. The neural progenitor-restricted isoform of the mark4 gene in 19q13. 2 is upregulated in human gliomas and overexpressed in a subset of glioblastoma cell lines. Oncogene, 22 (17):2581–2591, 2003.

6. Ivana Magnani, Chiara Novielli, Laura Fontana, Silvia Tabano, Davide Rovina, Ramona F Moroni, Dario Bauer, Stefania Mazzoleni, Elisa A Colombo, Gabriella Tedeschi, et al. Differential signature of the centrosomal mark4 isoforms in glioma. Analytical Cellular Pathology, 34(6):319–338, 2011.

7. Tatsushi Kato, Seij Satoh, Hiroshi Okabe, Osamu Kitahara, Kenji Ono, Chikashi Kihara, Toshihiro Tanaka, Tatsuhiko Tsunoda, Yoshio Yamaoka, Yusuke Nakamura, et al. Isolation of a novel human gene, markli, homologous to mark3 and its involvement in hepatocellular carcinogenesis. Neoplasia, 3(1):4–9, 2001.

8. Farha Naz, Mohd Shahbaaz, Krishna Bisetty, Asimul Islam, Faizan Ahmad, and Md Imtaiyaz Hassan. Designing new kinase inhibitor derivatives as therapeutics against common complex diseases: structural basis of microtubule affinity-regulating kinase 4 (mark4) inhibition. OMICS: A Journal of Integrative Biology, 19(11):700–711, 2015.

9. Huma Naz, Ehtesham Jameel, Nasimul Hoda, Ashutosh Shandilya, Parvez Khan, Asimul Islam, Faizan Ahmad, B Jayaram, and Md Imtaiyaz Hassan. Structure guided design of potential inhibitors of human calcium–calmodulin dependent protein kinase iv containing pyrimidine scaffold. Bioorganic & Medicinal Chemistry Letters, 26(3):782–788, 2016.

10. Sajjad Ahrari, Fatemeh Khosravi, Ali Osouli, Amirhossein Sakhteman, Alireza Nematollahi, Younes Ghasemi, and Amir Savardashtaki. Mark4 protein can explore the active-like conformations in its non-phosphorylated state. Scientific Reports, 9(1):12967, 2019.

11. Xianyan Shen, Xuesha Liu, Shunli Wan, Xin Fan, Huaiyu He, Rong Wei, Wenchen Pu, Yong Peng, and Chun Wang. Discovery of coumarin as microtubule affinity-regulating kinase 4 inhibitor that sensitize hepatocellular carcinoma to paclitaxel. Frontiers in Chemistry, 7:366, 2019.

12. Babita Aneja, Nashrah Sharif Khan, Parvez Khan, Aarfa Queen, Afzal Hussain, Md Tabish Rehman, Mohamed F Alajmi, Hesham R El-Seedi, Sher Ali, Md Imtaiyaz Hassan, et al. Design and development of isatin-triazole hydrazones as potential inhibitors of microtubule affinity-regulating kinase 4 for the therapeutic management of cell proliferation and metastasis. European Journal of Medicinal Chemistry, 163:840–852, 2019.

13. Mudasir Nabi Peerzada, Parvez Khan, Nashrah Sharif Khan, Fernando Avecilla, Shadab Miyan Siddiqui, Md Imtaiyaz Hassan, and Amir Azam. Design and development of small-molecule arylaldoxime/5-nitroimidazole hybrids as potent inhibitors of mark4: a promising approach for target-based cancer therapy. ACS omega, 5(36):22759–22771, 2020.

14. Bernhard Trinczek, Miro Brajenovic, Andreas Ebneth, and Gerard Drewes. Mark4 is a novel microtubule-associated proteins/microtubule affinity-regulating kinase that binds to the cellular microtubule network and to centrosomes. Journal of Biological Chemistry, 279(7): 5915–5923, 2004.

15. John S Sack, Mian Gao, Susan E Kiefer, Joseph E Myers, John A Newitt, Sophie Wu, and Chunhong Yan. Crystal structure of microtubule affinity-regulating kinase 4 catalytic domain in complex with a pyrazolopyrimidine inhibitor. Acta Crystallographica Section F: Structural Biology Communications, 72(2):129–134, 2016.

16. Narendran Annadurai, Khushboo Agrawal, Petr Džubák, Marián Hajdúch, and Viswanath Das. Microtubule affinity-regulating kinases are potential druggable targets for alzheimer’s disease. Cellular and molecular life sciences, 74:4159–4169, 2017.

17. Fangfang Li, Zongliang Liu, Heyuan Sun, Chunmei Li, Wenyan Wang, Liang Ye, Chunhong Yan, Jingwei Tian, and Hongbo Wang. Pcc0208017, a novel small-molecule inhibitor of mark3/mark4, suppresses glioma progression in vitro and in vivo. Acta Pharmaceutica Sinica B, 10(2):289–300, 2020.

18. Anas Shamsi, Saleha Anwar, Taj Mohammad, Mohamed F Alajmi, Afzal Hussain, Md Tabish Rehman, Gulam Mustafa Hasan, Asimul Islam, and Md Imtaiyaz Hassan. Mark4 inhibited by ache inhibitors, donepezil and rivastigmine tartrate: Insights into alzheimer’s disease therapy. Biomolecules, 10(5):789, 2020.

19. Ahmad Abu Turab Naqvi, Deeba Shamim Jairajpuri, Omar Mohammed Ali Noman, Afzal Hussain, Asimul Islam, Faizan Ahmad, Mohammed F Alajmi, and Md Imtaiyaz Hassan. Evaluation of pyrazolopyrimidine derivatives as microtubule affinity regulating kinase 4 inhibitors: Towards therapeutic management of alzheimer’s disease. Journal of Biomolecular Structure and Dynamics, 38(13):3892–3907, 2020.

20. Mudasir Nabi Peerzada, Parvez Khan, Nashrah Sharif Khan, Aysha Gaur, Fernando Avecilla, Md Imtaiyaz Hassan, and Amir Azam. Identification of morpholine based hydroxylamine analogues: selective inhibitors of mark4/par-1d causing cancer cell death through apoptosis. New Journal of Chemistry, 44(38):16626–16637, 2020.

21. Saleha Anwar, Shama Khan, Anas Shamsi, Farah Anjum, Alaa Shafie, Asimul Islam, Faizan Ahmad, and Md Imtaiyaz Hassan. Structure-based investigation of mark4 inhibitory potential of naringenin for therapeutic management of cancer and neurodegenerative diseases. Journal of Cellular Biochemistry, 122(10):1445–1459, 2021.

22. P Khan, S Rahman, A Queen, S Manzoor, F Naz, GM Hasan, S Luqman, J Kim, A Islam, F Ahmad, et al. Elucidation of dietary polyphenolics as potential inhibitor of microtubule affinity regulating kinase 4: In silico and.

23. Farha Naz, Faez Iqbal Khan, Taj Mohammad, Parvez Khan, Saaliqa Manzoor, Gulam Mustafa Hasan, Kevin A Lobb, Suaib Luqman, Asimul Islam, Faizan Ahmad, et al. Investigation of molecular mechanism of recognition between citral and mark4: A newer therapeutic approach to attenuate cancer cell progression. International journal of biological macromolecules, 107:2580–2589, 2018.

24. Parvez Khan, Aarfa Queen, Taj Mohammad, Smita, Nashrah Sharif Khan, Zubair Bin Hafeez, Md Imtaiyaz Hassan, and Sher Ali. Identification of α-mangostin as a potential inhibitor of microtubule affinity regulating kinase 4. Journal of natural products, 82(8):2252–2261, 2019.

25. Saleha Anwar, Anas Shamsi, Rajiv K Kar, Aarfa Queen, Asimul Islam, Faizan Ahmad, and Md Imtaiyaz Hassan. Structural and biochemical investigation of mark4 inhibitory potential of cholic acid: Towards therapeutic implications in neurodegenerative diseases. International Journal of Biological Macromolecules, 161:596–604, 2020.

26. J Ferlay, M Ervik, F Lam, M Colombet, L Mery, M Piñeros, A Znaor, I Soerjomataram, and F Bray. Global cancer observatory: Cancer today. lyon fr. int. agency res. Cancer, 3:2019, 2018.

27. Hyuna Sung, Jacques Ferlay, Rebecca L. Siegel, Mathieu Laversanne, Isabelle Soerjomataram, Ahmedin Jemal, and Freddie Bray. Global cancer statistics 2020: Globocan estimates of incidence and mortality worldwide for 36 cancers in 185 countries. CA: A Cancer Journal for Clinicians, 71(3):209–249, 2021. doi: https://doi.org/10.3322/caac.21660.

28. Gordon M Cragg and David J Newman. Plants as a source of anti-cancer agents. Journal of ethnopharmacology, 100(1-2):72–79, 2005.

29. Md Rashedur Rahman, Shammi Binte Bashar, Rakibul Hasan Rifat, Md Shah Poran, Md Ashikur Rahman, Fargana Islam, and Bidhan Chandro Saha. Medicinal plants with anticancer effects available in bangladesh: A review. Journal of Pharmacognosy and Phytochemistry, 10(3):41–49, 2021.

30. Riki Nova, Aida SD Hoemardani, and Melva Louisa. Potential of herbal medicines in cancer therapy. The Indonesian Journal of Cancer Control, 1(1):32–42, 2021.

31. JG Graham, ML Quinn, DS Fabricant, and NR Farnsworth. Plants used against cancer–an extension of the work of jonathan hartwell. Journal of ethnopharmacology, 73(3):347–377, 2000.

32. Anchala I Kuruppu, Priyani Paranagama, and Charitha L Goonasekara. Medicinal plants commonly used against cancer in traditional medicine formulae in sri lanka. Saudi Pharmaceutical Journal, 27(4):565–573, 2019.

33. Ghulam Mustafa, Rawaba Arif, Asia Atta, Sumaira Sharif, Amer Jamil, et al. Bioactive compounds from medicinal plants and their importance in drug discovery in pakistan. Matrix Sci. Pharma, 1(1):17–26, 2017.

34. Thomas A Halgren, Robert B Murphy, Richard A Friesner, Hege S Beard, Leah L Frye, W Thomas Pollard, and Jay L Banks. Glide: a new approach for rapid, accurate docking and scoring. 2. enrichment factors in database screening. Journal of medicinal chemistry, 47(7):1750–1759, 2004.

35. Schrödinger SoftwareRelease 2015-2. QikProp 4.4 user manual, 2015.

36. Schrödinger Release. 1: Desmond molecular dynamics system, de shaw research, new york, ny, 2019. maestro-desmond interoperability tools, schrödinger, new york, ny, 2019, 2019.

37. Khan AM Kothapalli R, Chong YS Basappa, Gopalsamy A, and Annamalai L. Cheminformatics-based drug design approach for identification of inhibitors targeting the characteristic residues of mmp-13 hemopexin domain. PloS one, 2010.

38. Antonija Kuzmanic and Bojan Zagrovic. Determination of ensemble-average pairwise root mean-square deviation from experimental b-factors. Biophysical journal, 98(5):861–871, 2010.

39. Zeeshan Ahmed Sheikh, Saleha Suleman Khan Aqib Zahoor, and Khan Usmanghani. Design, development and phytochemical evaluation of a poly herbal formulation linkus syrup. Chinese Medicine, 5(2), 2014.

40. Tran Thi Minha, Nguyen Thi Hoang Anh, Vu Dao Thanga, and Tran Van Sung. Study on chemical constituents of salacia chinensis l. collected in vietnam. Zeitschrift für Naturforschung B, 63(12):1411–1414, 2008.

